# Choice of assemblers has a critical impact on de novo assembly of SARS-CoV-2 genome and characterizing variants

**DOI:** 10.1101/2020.12.15.422939

**Authors:** Rashedul Islam, Rajan Saha Raju, Nazia Tasnim, Md. Istiak Hossain Shihab, Maruf Ahmed Bhuiyan, Yusha Araf, Tofazzal Islam

## Abstract

**Background:** Coronavirus Disease 2019 (COVID-19), caused by severe acute respiratory syndrome coronavirus 2 (SARS-CoV-2) has become a global pandemic following its initial emergence in China. Using next-generation sequencing technologies, a large number of SARS-CoV-2 genomes are being sequenced at an unprecedented rate and being deposited in public repositories. For the de novo assembly of the SARS-CoV-2 genomes, a myriad of assemblers is being used, although their impact on the assembly quality has not been characterized for this virus. In this study, we aim to understand the variabilities on assembly qualities due to the choice of the assemblers.

**Results:** We performed 6,648 *de novo* assemblies of 416 SARS-CoV-2 samples using 8 different assemblers with different k-mers. We used Illumina paired-end sequencing reads and compared the genome assembly quality to that of different assemblers. We showed the choice of assemblers plays a significant role in reconstructing the SARS-CoV-2 genome. Two metagenomic assemblers e.g. MEGAHIT and metaSPAdes performed better compared to others in most of the assembly quality metrics including, recovery of a larger fraction of the genome, constructing larger contigs and higher N50, NA50 values etc. We showed that at least 09% (259/2,873) of the variants present in the assemblies between MEGAHIT and metaSPAdes are unique to the assembly methods.

**Conclusion:** Our analyses indicate the critical role of assembly methods for assembling SARS-CoV-2 genome using short reads and their impact on variant characterization. This study could help guide future studies to determine which assembler is best suited for the de novo assembly of virus genomes.

## Introduction

SARS-CoV-2 is the seventh member of the Coronaviridae family to infect humans, which is responsible for the current COVID-19 pandemic (Zhu *et al.*, 2020). This virus is ravaging the world with more than 1.5 million deaths in the year 2020. To understand its pathophysiological mechanism, mutation pattern, epidemiological tracing and transmission pathways, the single-stranded RNA genome of SARS-CoV-2 has been sequenced in different countries. Around 50,000 SARS-CoV-2 genomes (both complete or partial) have been submitted to NCBI Nucleotide records and Nextstrain database since the first whole-genome was sequenced in January 2020 *(Hadfield et al., 2018; Wu et al., 2020)*. This unprecedented speed of genome sequencing was possible due to the advancement in sequencing technologies and the availability of open-source bioinformatics tools.

By the end of 2020, 84% (140,837/168,547) of SARS-CoV-2 sequencing runs deposited on NCBI’s Sequence Read Archive were generated using Illumina short-read sequencing technology (Leinonen, Sugawara and Shumway, 2011). Along with short-read sequencing technologies, long-read sequencing technologies were also used in combination with short reads or alone to decipher SARS-CoV-2 genome (Lu *et al.*, 2020). Many assembly tools (assemblers) are publicly available to assemble the genome from short reads. These assemblers use a combination of, or solely, these methods: De Bruijn graph, Overlay Layout Consensus and greedy graph method. The quality of the virus genome assembly varies depending on the assembler of choice, genome composition, depth of sequencing, sample preparation etc (Sutton *et al.*, 2019). Due to the rapid pace of SARS-CoV-2 genome sequencing, use of different sequencing assays and availability of multiple assemblers, the assemblers need to be benchmarked and updated for the de novo assembly of the SARS-CoV-2 genome.

RNA viruses naturally accumulate random genetic variations during the course of infection (Sanjuán and Domingo-Calap, 2016). Mutations in the genomes are used to track the transmission of SARS-CoV-2 virus in which closely related genomes are anticipated to be closely related infections (Hadfield *et al.*, 2018). Phylogeny of genomes is used to cluster similar clades where the genetic diversity solely depends on the variants present on the genomes (Volz, Koelle and Bedford, 2013). However, assemblers could introduce erroneous base(s) in the consensus sequences due to their error correction method, quality filtering or parameter selections *(Baker, 2012; Swain et al., 2012; Olson et al., 2015)*. Assemblers have their unique error profiles and therefore genomic variants also vary by assemblers (Salzberg *et al.*, 2012). Here we sought to find out the instances of genomic variants that were solely driven by the choice of assemblers.

To date, the degree of variation in assembly qualities among different assemblers has not been reported for SARS-CoV-2. In this study, we present a comprehensive investigation on the performance of assemblers for SARS-CoV-2 genome assembly with publicly available Illumina paired-end datasets. We compared how different attributes affect the performance of these assemblers in terms of percentage of genome recovery, largest contig, total length, N50, NA50, L50, LA50 etc. We investigated the variants on the assembled contigs and showed variant detection varies between assemblers.

## Results

### De novo assemblers showed marked differences in assembly quality

Illumina paired-end data for the SARS-CoV-2 genomes have been collected from different assay types e.g., amplicon, WGA, RNA-Seq, targeted-capture and other **(Fig. 1a)**. We randomly selected 100 libraries for each assay type. In our pipeline, some samples did not finish the job and were removed from subsequent analysis **(Fig. 1b, Table 1)**. To evaluate assembly quality among different assemblers in different assay types, we selected four *de novo* metagenome assemblers (e.g., metaSPAdes, MetaVelvet, MEGAHIT and Ray Meta) and four *de novo* genome/transcriptome assemblers (e.g., AbySS, velvet, SPAdes and Trinity) **(Table S1)**. To rule out the influence of the choice of k-mers on assembly quality we used three k-mers (21, 63 and 99) for the assemblers requiring a fixed k-mer to run.

**Figure 1:**
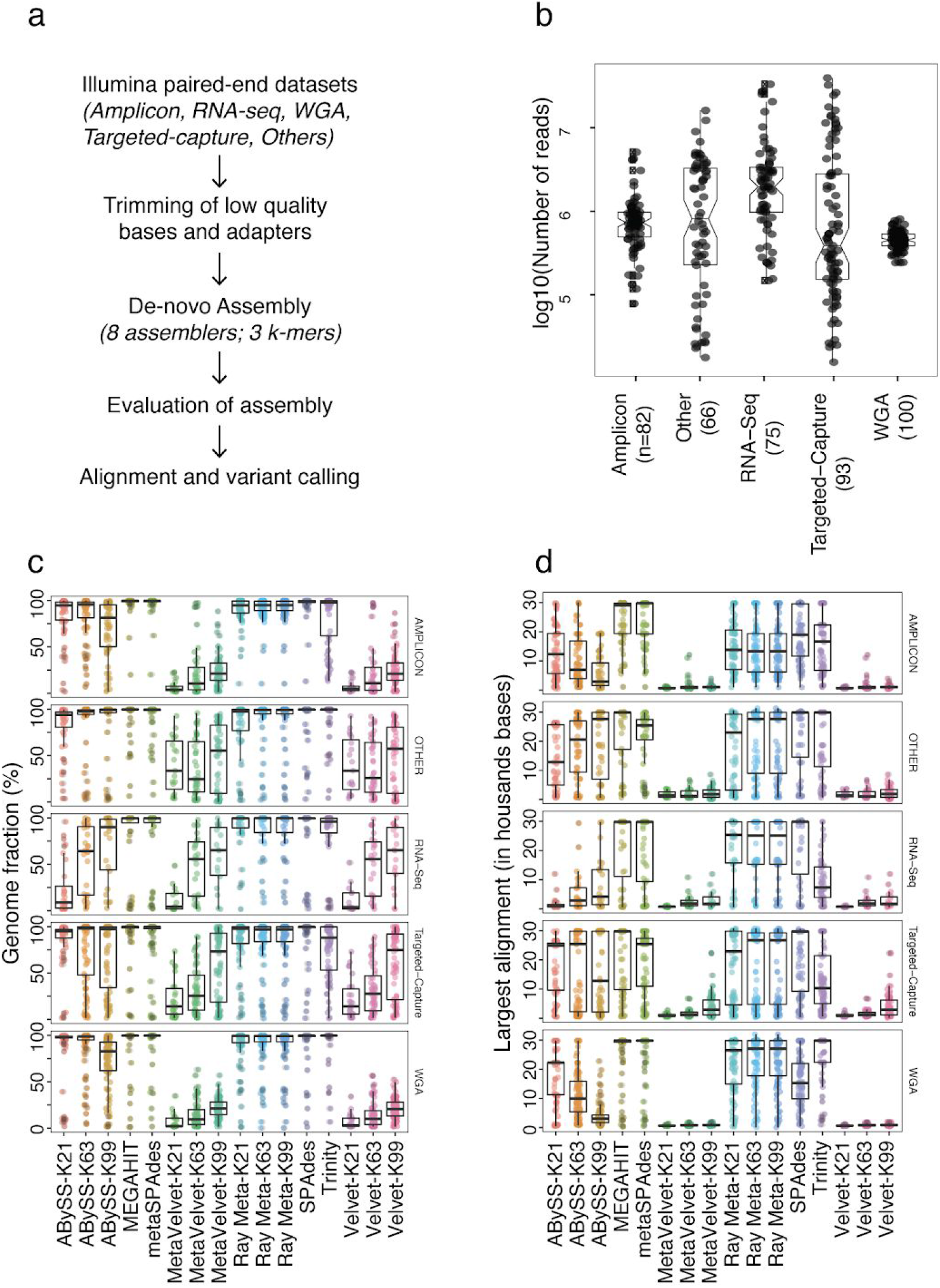
Comparison of assemblers for different sequencing assays. **a)** Experimental strategies to assemble SARS-CoV-2 genomes using different assemblers and calling variants from the assembled contigs. From different sequencing assay types, samples were randomly selected for assembly and subsequent analysis. **b)** Total number of reads for each sample across different assay types. **c)** Fraction of SARS-CoV-2 genome assembled by different assemblers. **d)** The largest continuous alignment in the assembly produced by different assemblers.

**Table 1:** Summary of the viral RNA dataset of SARS-CoV-2 available by 14th June 2020

| Assay type | Library source | Library layout | # libraries | # selected |
| --- | --- | --- | --- | --- |
| AMPLICON | VIRAL RNA | PAIRED | 5254 | 82 |
| OTHER | VIRAL RNA | PAIRED | 68 | 66 |
| RNA-Seq | VIRAL RNA | PAIRED | 682 | 75 |
| Targeted-Capture | VIRAL RNA | PAIRED | 194 | 93 |
| WGA | VIRAL RNA | PAIRED | 1061 | 100 |

Genome fraction recovery was highly variable across the different assemblers **(assembly quality terminologies; Table S2)**. Most of the assemblers (e.g., ABySS, Ray Meta, SPAdes, Trinity) recovered a larger fraction (median of genome fraction >90%) of the genome whereas MEGAHIT and metaSPAdes recovered almost the entire genome (median of >= 99.7%) across all assay types. Velvet and Metavelvet recovered a lower fraction of the genome compared to other assemblers invariably across assay types, despite an increase in genome fraction recovery with increasing k-mer **(Fig. 1c)**. We confirmed that the recovery of the fraction of the genome by different assemblers was not affected by the sequencing depth of the libraries in different assay types **(Fig. 2a)**. For all assemblers, genome fractions were invariably recovered across the span of the sequencing depth except for Targeted-Capture assay which showed a negative correlation (Spearman: −0.24). Consistent with the recovery of the genome, the larger uninterrupted alignment of assemblies to the reference genome were obtained by MEGAHIT (median of >29,000bp) and metaSPAdes (median of >25,000bp) across all assay types **(Fig. 1d)**. In contrast, Velvet and MetaVelvet showed relatively lower contiguous alignments. MEGAHIT and metaSPAdes also generated a higher number of assemblies with 90 percent of the genome covered by a single contig whereas ABySS was unable to generate longer contiguous sequences at different k-mers **(Fig. 2b)**.

**Figure 2:**
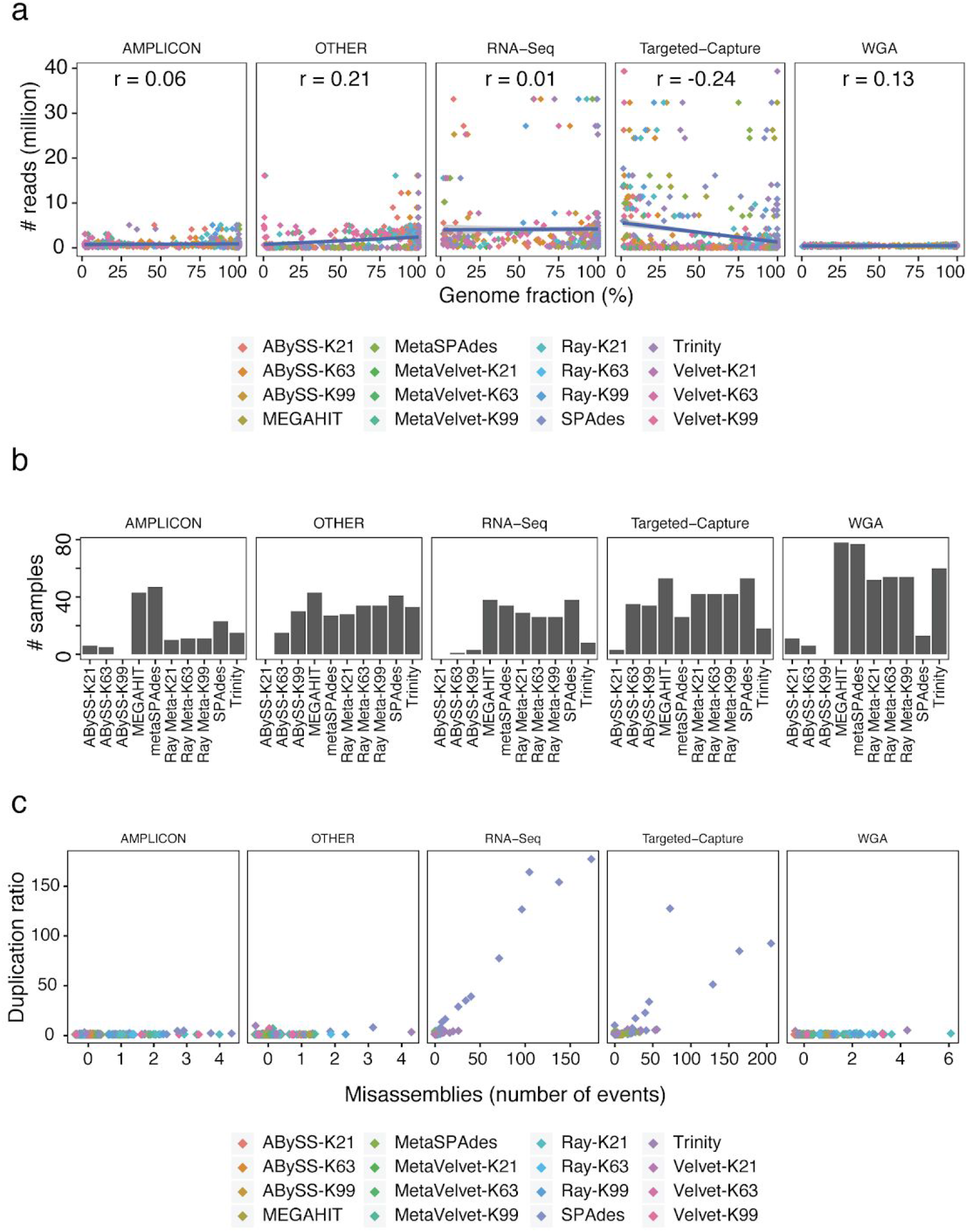
Assessment of sequencing and assembly quality. **a)** Pearson correlation coefficient between the number of reads and fraction of genome assembled in respect to assay types. **b)** Assemblies with 90 percent of the genome covered by a single contig. **c)** Relation between misassemblies and duplication ratio across assay types for all the assemblers. Random jitter was added to avoid overplotting of points on the discrete x-axis values.

Number of misassemblies and duplication ratio were low in different assays except for RNA-seq and Targeted-Capture. On average, SPAdes produced the higher number of misassembled events along with higher duplication ratio **(Fig. 2c, Fig. S1a)**. However, 87% (4,291/4,912) of the assemblies did not produce any misassembled events. ABySS, Velvet and MetaVelvet produced the minimum number of misassemblies across different k-mers and assay types. The duplication ratio was higher in SPAdes and Trinity compared to other assemblers **(Fig. S1b)**. The duplication ratio varies with increased k-mers in ABySS, Velvet, MetaVelvet, Ray Meta. The median duplication ratio for all assemblies is 1.005 where 25% (1,247/4,912) of the assemblies had duplication ratio equal to 1.

### MEGAHIT and metaSPAdes performed better in most of the assembly quality matrices

N50 is the minimum contig length needed to construct 50% of the genome. There were large amounts of variations between assemblers for N50. All assemblers performed poorly on the RNA-seq assay type **(Fig. 3a)**. For the rest of the assay types, MEGAHIT, metaSPAdes, Ray Meta, SPAdes and Trinity performed better while ABySS, Velvet and MetaVelvet showed larger variations in N50 depending on the assay types and k-mers. NA50 is an improved matrix of the N50 contig length which breaks contigs into aligned blocks at misassembly events and removes all unaligned bases. MEGAHIT and metaSPAdes were able to generate larger NA50 values in all assay types compared to other assemblers **(Fig. 3b)**. Metavelvet and Velvet were unable to produce large N50 as well as NA50 values across all the assay types. MEGAHIT and metaSPAdes produced lower L50 (median = 1) and LA50 (median = 1) values compared to other assemblers across different assay types **(Fig. 3c,d)**. We have observed MEGAHIT and metaSPAdes outperformed other assemblers in several other quality matrices e.g., contig lengths, number of unaligned contigs, number of Ns, percent overlap with genes etc. **(Supplementary file 1,2)**.

**Figure 3:**
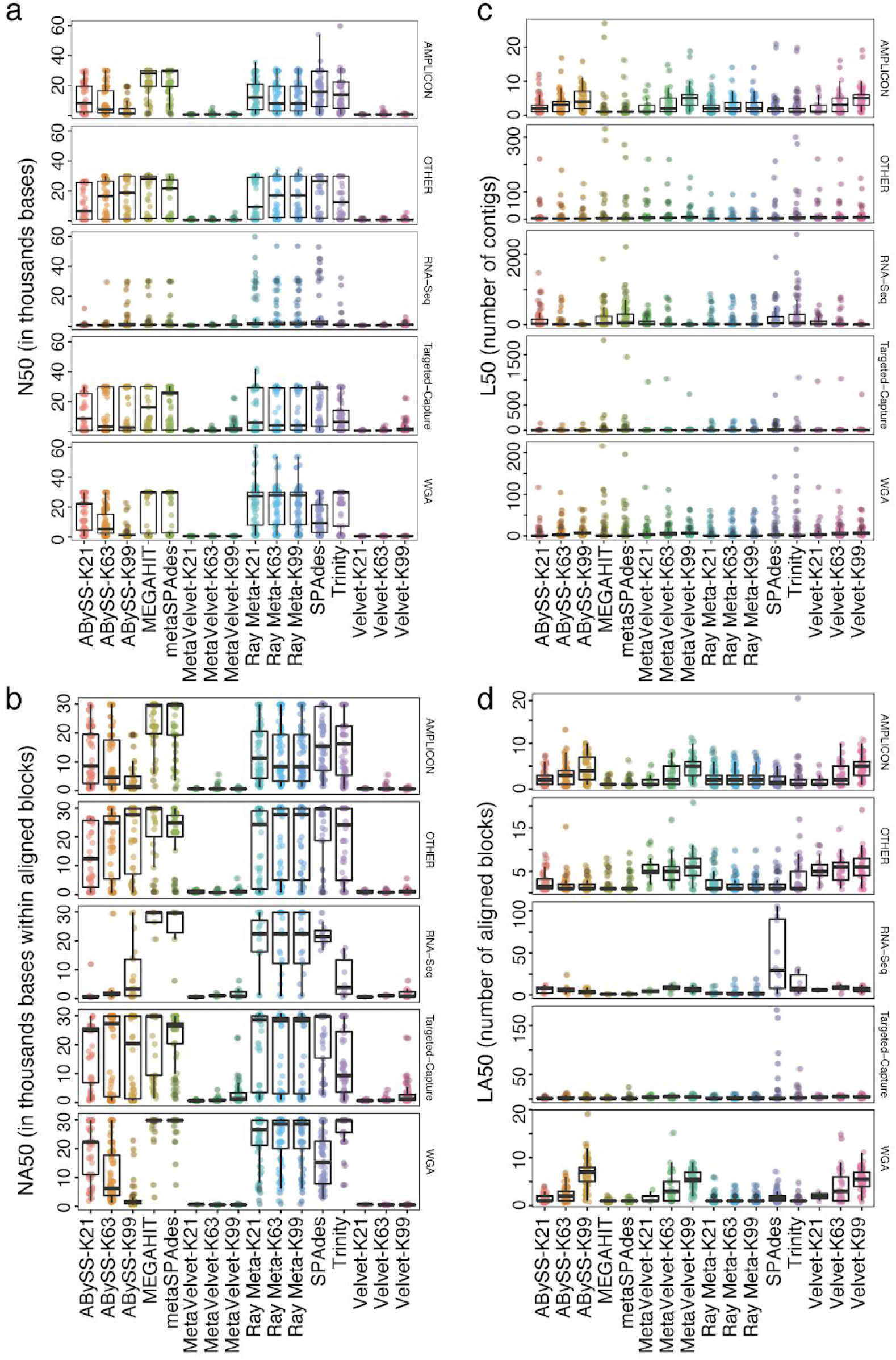
Comparison of contigs and aligned genomic blocks. **a,b)** Length of the smallest contig (N50) and aligned block (NA50) at 50% of the total genome length. **c,d)** Minimum number of contigs (L50) and aligned blocks (LA50) at 50% of the total genome length.

### SARS-CoV-2 genome assembly contiguity at the repeat region

We investigated the 585bp repeat region in the SARS-CoV-2 genome located at the 3’-end (MN908947.3:29870-29903). Most of the assemblers failed to assemble the repeat region and more 100bp gap was created at the 3’-end in different assemblies. To identify the presence of similar assembly gaps in the assemblies, we binned the entire genome into 50bp non-overlapping windows and counted the number of bases assembled in each bin throughout the genome **(Fig. 4a)**. We compared the assemblies of top-performing four assemblers e.g., MEGAHIT, metaSPAdes, Trinity and ABySS-K63 for 392 samples of five assay types. Gaps in the assemblies are shown in red color and assembled regions are in gray in the heatmap. We found that in addition to gaps in the 3’-end, there were assembly gaps at the 5’-end of the genome across the assemblies. On average there was a 50bp assembly gap at the 5’-end. Besides gaps in the 5’ and 3’-ends, we did not observe consistent assembly gaps in the SARS-CoV-2 genome.

**Figure 4:**
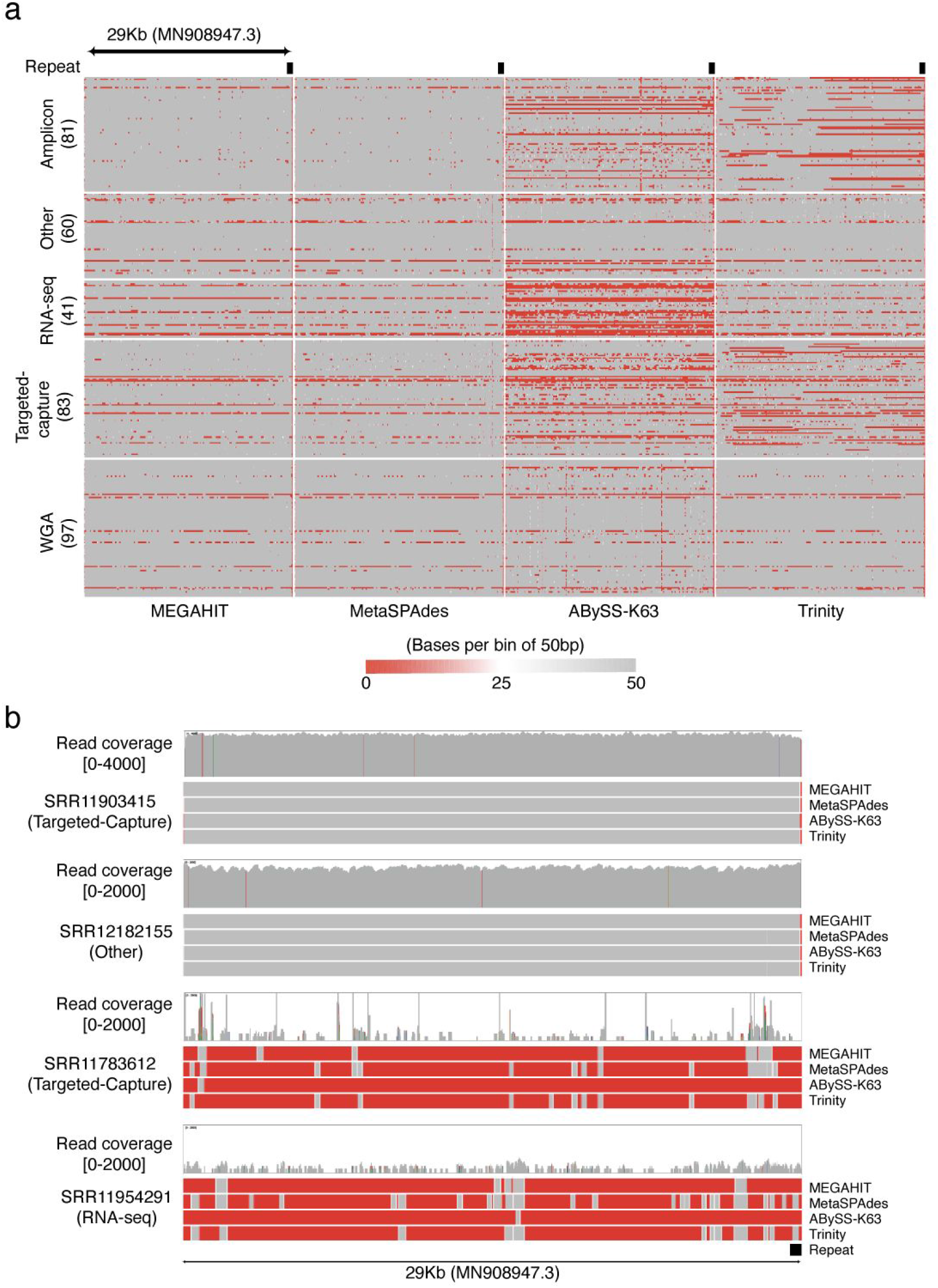
Analysis of assembly contiguity and gap. **a)** SARS-CoV-2 genome was binned into 50bp non-overlapping windows. For each bin, number of bases assembled were plotted in the heatmap. Using continuous color scale, genomic bins were plotted by the number of bases assembled at those bins where 50bp assembled bins shown in gray and 0bp assembled bins were in red color. Samples with successful assemblies for all four assemblers were included here. **b)** Two samples (SRR11903415 and SRR12182155) with contiguous assembly (gray) and two samples (SRR11783612 and SRR11954291) with gapped assembly (red) were plotted with sequencing read coverage across the SARS-CoV-2 genome.

Independent of assemblers, some samples showed better contiguity in assembly or worse quality assembly with numerous gaps. To investigate the sample-specific assembly quality variations, we analyzed four samples consisting of two better and two worse quality assemblies. The samples (SRR11903415 and SRR12182155) with better quality assembly had uniform read coverage throughout the genome whereas other two samples (SRR11783612 and SRR11954291) showed gapped assembly independent of assemblers due to lack of read coverage at the gapped regions. We visualized the read densities in the genome along with the gaps (red) and assembled regions (grey) **(Fig. 4b)**. This suggests that without uniform read distributions throughout the genome and sufficient coverage assemblers fail.

### Influence of k-mer on assembly quality

All 8 assemblers we tested use graph-based methods where the choice of k-mer size affects the contiguity of an assembly (Chikhi and Medvedev, 2014; Vollmers, Wiegand and Kaster, 2017). Four of the *de novo* assemblers (e.g., ABySS, Velvet, MetaVelvet and Ray Meta) we used require a single k-mer. To analyze the variability in assembly quality across 416 samples, we used three different k-mers (i.e., 21, 63 and 99). With different k-mers ABySS, Velvet and MetaVelvet showed variabilities in assembly quality but Ray Meta did not show larger variabilities on average **(Fig. S2)**. ABySS performed relatively better with smaller k-mers in recovering fraction of genome, N50, NA50 values with lower misassemblies and duplication ratio. Velvet and MetaVelvet performed better with higher k-mers at the cost of higher misassemblies and duplication ratios. However, L50 values improved overall with the increasing k-mer values for all four assemblers.

### Variant calling from de novo assemblies varies between assemblers

We aligned the assembled contigs to the reference genome and called the variants **(see Methods)**. For 416 samples, the assemblers produced different numbers of variants which are expectedly correlated (Pearson = 0.73) with the fraction of genomes assembled by different assemblers **(Fig. S3)**. To investigate the occurrences of variants exclusively present in a given assembler, we compared the variants identified from the two best-performing assemblers e.g., MEGAHIT and metaSPAdes. We took the common genomic regions assembled by both MEGAHIT and metaSPAdes for all samples. We found 25% (95/385) of the samples had at least one assembler specific variant. Among all the variants identified in the common assembly regions, 92% of the variants overlapped between two assemblers **(Fig. 5a**). To further understand the consequence of assembler specific variants in biologically important genomic features, we analyzed the Spike (S) gene locus and found that 06% (23/385) of the samples have assembler specific variants in the Spike locus **(Fig. 5b)**. Here we showed the example of variants in the Spike locus which are concordant (ERR4208998) between MEGAHIT and metaSPAdes, discordant (SRR11783589) and has different variant due to assembly gap (SRR12182180) in MEGAHIT. In many instances raw reads do not contain those variants and suggesting that spurious variants arise due to assembly errors **(Fig. S3b, S4a,b)**. This highlights the importance of correcting assembler specific spurious variants before functional characterization of the variants and analysing pan-genomes using *de novo* assemblies for phylogeny construction.

**Figure 5:**
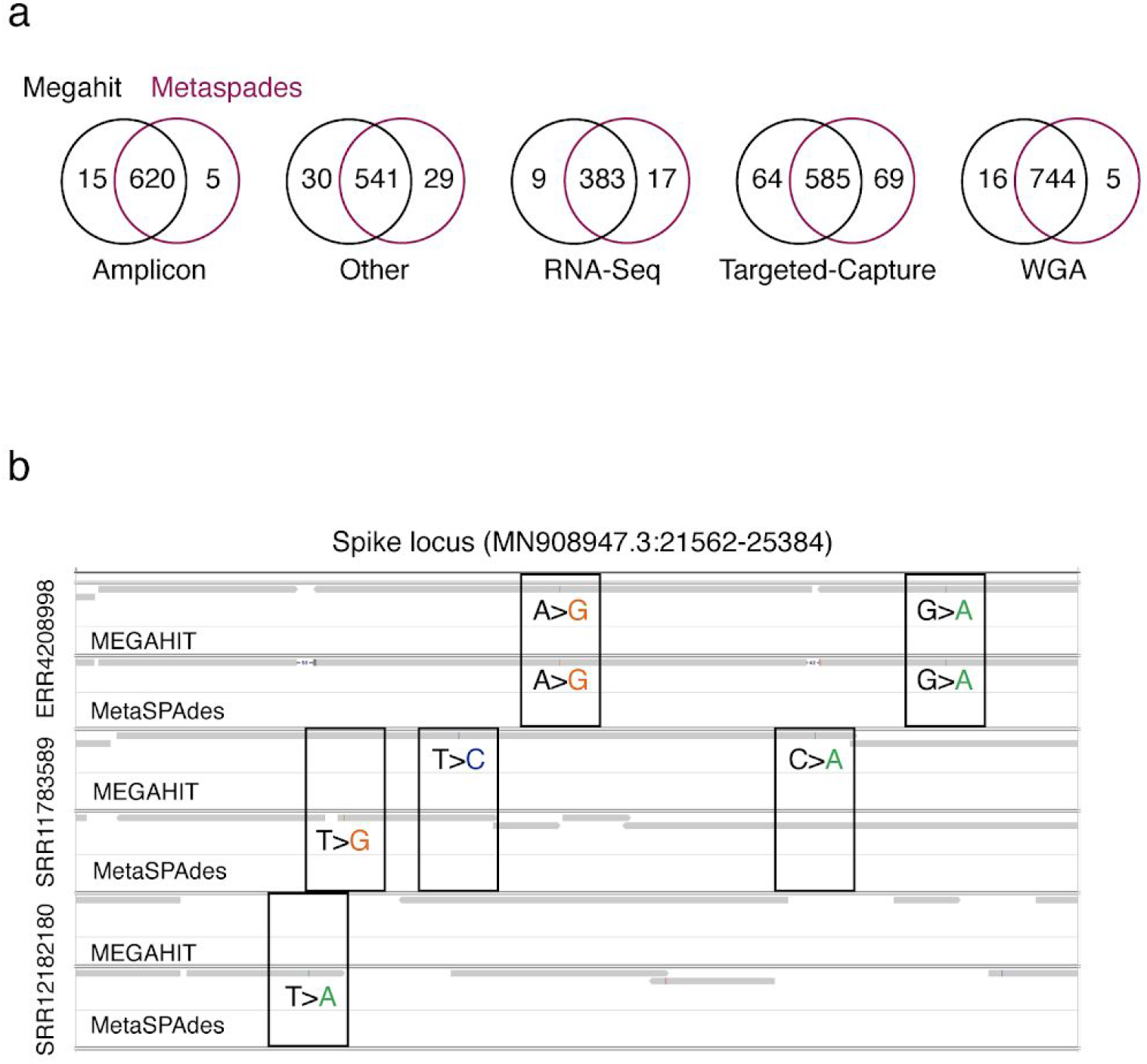
Differences in genomic variants between two assemblers. a) Overlap of variants within the same genomic blocks common between MEGAHIT and metaSPAdes assemblies for different assay types. Single nucleotide variants and short insertions/deletions are included here. b) Example of concordant (ERR4208998) and discordant (SRR11783589, SRR12182180) variants between MEGAHIT and metaSPAdes. Contigs are represented in gray bars and variant nucleotides are highlighted.

**Figure 6:**
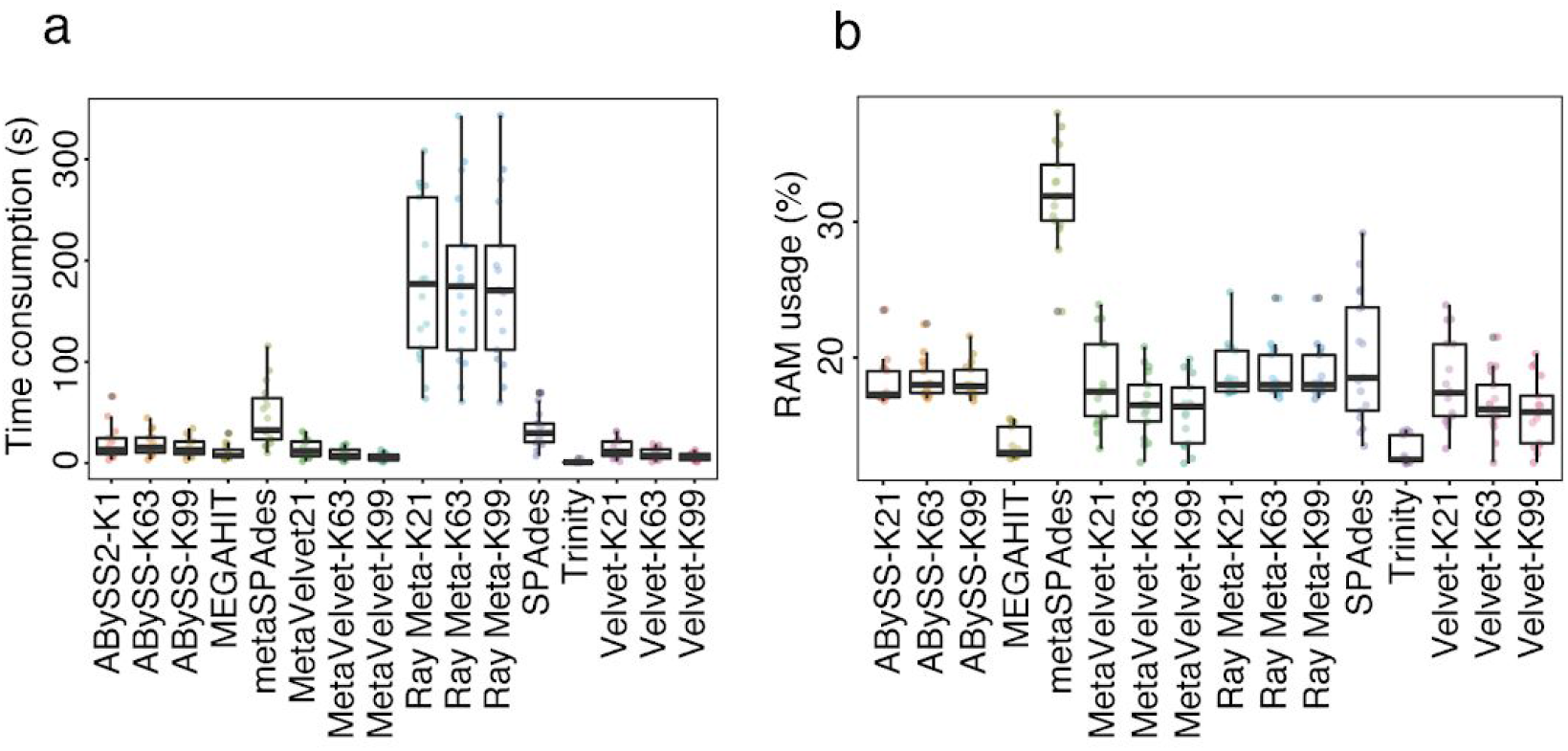
Computational resources required for different assemblers. a) Time consumed by different assemblers in CPU seconds. b) RAM percentage usage by different assemblers.

### Computational performance by different assemblers

Scarcity of computational resources also forces us to pay special attention to space and time complexity during *de novo* assembly of genomes. For calculating the Central Processing Unit (CPU) time consumption and Random Access Memory (RAM) usage, we randomly selected 17 amplicon libraries with around one million paired-end reads. We utilized 4 cores and 8 threads for all assemblers on a dedicated computer. Assembly completion time has been adopted as time consumed in CPU seconds **(Fig. 6a)**. Trinity and MetaVelvet-K99 consumed the lowest time among all assemblers with median values of 0.73 and 4.1 seconds respectively (**Fig. 6a**). Ray Meta and metaSPAdes took the longest time. With regards to RAM usage, we observed that Trinity and MEGAHIT required the least amount of RAM with median values of 12.5% and 13.0% respectively (**Fig. 6b**). metaSPAdes required the highest amount (31.9%) of RAM compared to other assemblers. Notably, the choice of k-mer for the same assembler had little to no effect on the CPU time and RAM usage measurements.

## Discussion

In this study, we compared 16 assembler variations using eight de novo assemblers for the benchmarking of the assembly quality in SARS-CoV-2. We observed two metagenomic assemblers e.g., MEGAHIT and metaSPAdes outperformed other assemblers in regards to the genome fraction recovery, largest contig length, N50 length, NA50 length, L50 and LA50 contig number. The fraction of genome recovery could be 10-folds different between assemblers e.g., MEGAHIT (99%) vs MetaVelvet-K21 (10%). Although all 8 assemblers used the de Bruijn graph method for de novo assembly, the differences we observed are due to the variations in their implementation, error correction, quality thresholds and choice of other parameters. Despite better performances by the metagenomic assemblers the entire viral genome was not assembled in most cases, especially at the termini of the genome. Therefore, there is a need to develop newer assembly methods specially designed to assemble complete viral genomes.

Single nucleotide variants and short insertions and deletions vary by the assemblers, possibly correlated to the “aggressiveness” of the assembler (Salzberg *et al.*, 2012). Differences in variants introduced by assemblers may have an impact on downstream comparative genomic applications, such as pan-genome comparison or constructing phylogenetic tree using de novo genome assemblies. Often assembler specific variants are the result of assembly errors. Such assembly errors can be resolved by using post-assembly genome improvement pipelines that use local assembly and/or raw read alignment to the erroneous variants (Swain *et al.*, 2012).

To discover novel viruses, the sequence of complete viral genomes is inevitable rather than fragmented viral contigs. Low read coverage and genomic repeats resulted in assemblies with poor genome recovery independent of assemblers. Recent benchmarking studies reported metagenomic assemblers resulted in the relatively higher contiguous viral assemblies using viral metagenomic data (Roux *et al.*, 2017; Sutton *et al.*, 2019). Our analysis for SARS-CoV-2 data, using different sequencing assay types, identified two metagenome assemblers e.g., MEGAHIT and MetaSPAdes addressed the challenges of virome data better than other assemblers. Our benchmarking data for SARS-CoV-2 genome can be used to choose suitable de novo assemblers for similar genomes.

## Methods

### Data source and annotation

Publicly available raw sequencing data of SARS-CoV-2 genome were acquired from Sequence Read Archive (SRA). In this study, we used paired-end Illumina sequencing libraries of viral RNA. We randomly selected 100 paired-end Illumina libraries from six different assay types e.g., amplicon, RNA-seq, targeted-capture, whole genome amplification (WGA) and other categories **(Table 1, Fig. 1b)**. As a reference of SARS-CoV-2, *‘MN908947.3’* genome version was used. For annotation of genomics features ‘Sars_cov_2.ASM985889v3.100.gff3’ was downloaded from the Ensembl database.

### Read pre-processing

Adapter and low-quality bases were trimmed using Trimmomatic (Bolger, Lohse and Usadel, 2014) with default parameters. Raw reads were quality checked using FastQC and multiQC (Andrews S., 2010; Ewels *et al.*, 2016) **(Fig. 1a)**.

### Assemblers tested

In this study, de novo assembly of paired-end reads was performed using the current versions of eight different short-read assemblers. We used ABySS assembler which is optimized for short reads. The parallel version of ABySS is capable of assembling large genomes (Simpson *et al.*, 2009). MEGAHIT is an ultra-fast and memory-efficient short-read assembler, optimized for metagenomes, also works well on generic single genome assembly of small or mammalian size (Li *et al.*, 2015). Ray Meta is used for metagenome assembly and profiling (Boisvert *et al.*, 2012). SPAdes can assemble sequences from single-cell and multicell data types (Bankevich *et al.*, 2012). The Velvet assembler was designed for short-read sequencing data (Zerbino and Birney, 2008). metaSPAdes is a metagenomic assembler and MetaVelvet is an extension of Velvet for metagenome assembly from short sequence reads (Namiki *et al.*, 2012; Nurk *et al.*, 2017). Trinity performs de novo transcriptome assembly (Grabherr *et al.*, 2011). For every assembler mentioned above, we have used default parameters unless otherwise mentioned. Fixed k-mer values 21, 63, 99 were used for ABySS, Velvet, MetaVelvet and Ray Meta. For the rest of the assemblers default k-mer value was applied **(Fig. 1a, Table S1)**.

### Generation of assembly quality matrix

To generate an assembly quality matrix using metaQUAST (Mikheenko, Saveliev and Gurevich, 2016), we removed the contigs with < 500bp and compared all the assemblies to SARS-CoV-2 *‘MN908947.3’* reference genome.

### Alignment

We aligned the assembled contigs to the SARS-CoV-2 *‘MN908947.3’* reference genome using Minimap2 (version 2.17 r941) (Li, 2018). In Minimap2, we used 5% divergence between reference and assembly sequences to ensure inclusion. To align Illumina paired-end reads, we used BWA tools (version 0.7.17 r1188) (Li, 2013) with its mem feature enabled and default parameters.

### Variant calling from assembled contigs

After aligning the assembled contigs to the reference genome using Minimap2, we sorted the contigs by coordinates and indexed using SAMtools (version 1.11)(Li *et al.*, 2009). BCFtools (version 1.1) mpileup utility was used to generate genotype likelihoods at each genomic position with coverage, from the sorted BAM files to raw VCF formats. To extract the variant calls from the VCF file, we used BCFtools’ ‘call’ command, with the default definition of the ‘--ploidy’ parameter. Other parameters were also unchanged, as suggested.

### CPU and RAM usage

To check the computational performance of the assemblers, we randomly took 17 samples from our dataset and used 4 cores (8 threads) to perform assembly. Time to accomplish the assembly by an assembler has been adopted as CPU time for the particular assembler. Mathematically,

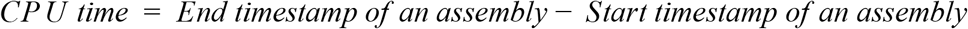

For finding RAM usage, we tracked the percentage usage of RAM every 0.5 seconds during assembly. We used a dedicated computer with 8 GB of RAM and accepted the maximum RAM usage among all values as final RAM usage. Mathematically,

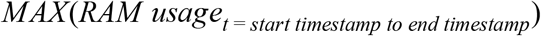

## Acknowledgements

We thank the authors of all datasets used in this study for the availability of their data.

## Author Contributions

Conceptualization, R.I.; Methodology, R.I., R.S.R., N.T. and M.I.H.S; Software, R.S.R., N.T., M. I.H.S and R.I.; Formal analysis, R.S.R., N.T., M.I.H.S, and R.I.; Investigation, R.I., R.S.R., N. T. and M.I.H.S; Writing - Original Draft, R.I., M.A.B., R.S.R., N.T., M.I.H.S and Y.A.; Writing - Review and Editing, R.I., M.A.B., Y.A. and T.I.; Visualization, R.I.; Supervision, R.I.; Project administration, Y.A.

## Supplementary information

**Figure S1.**
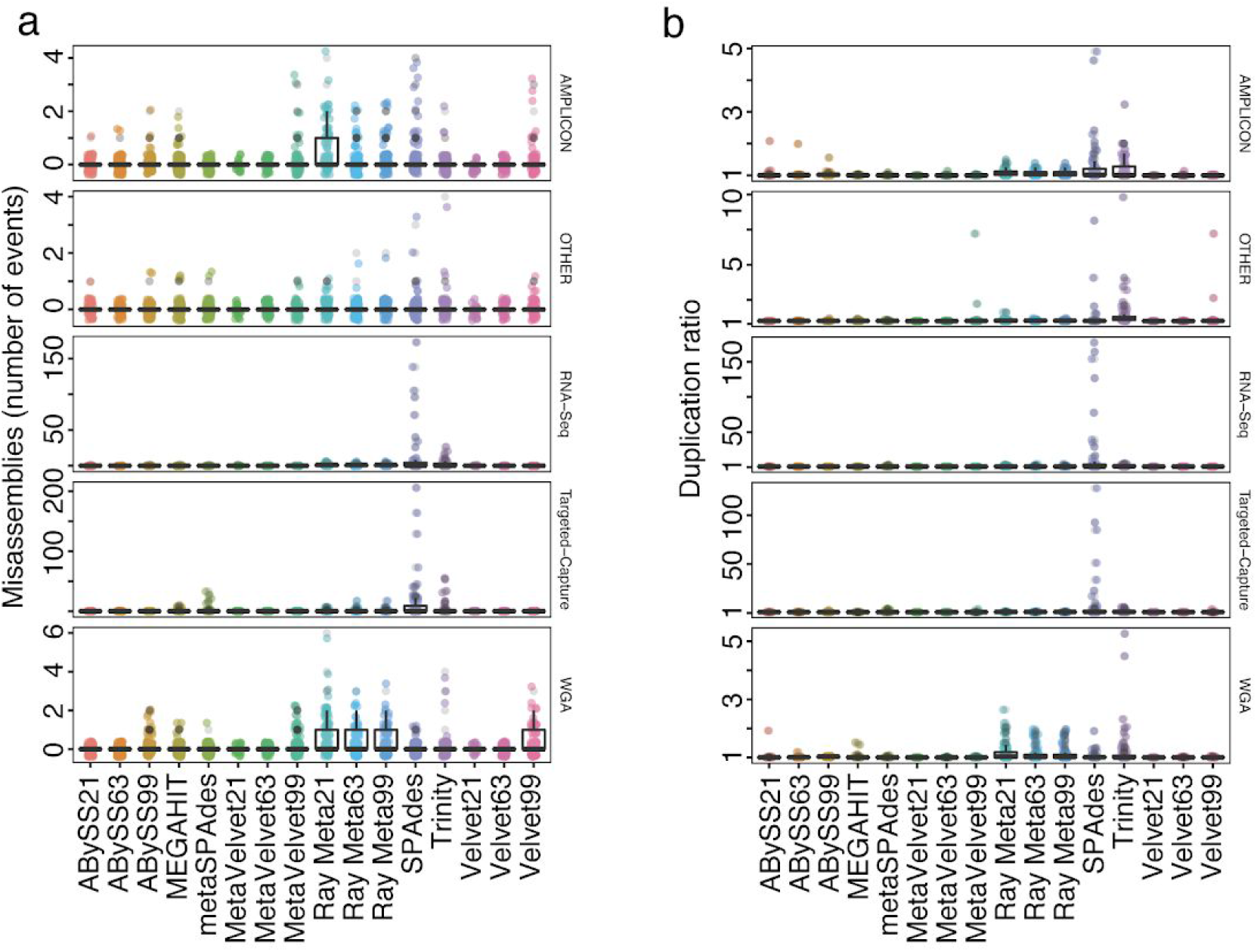
**a)** Number of events for misassemblies. **b)** The duplication ratios across different assays types.

**Figure S2.**
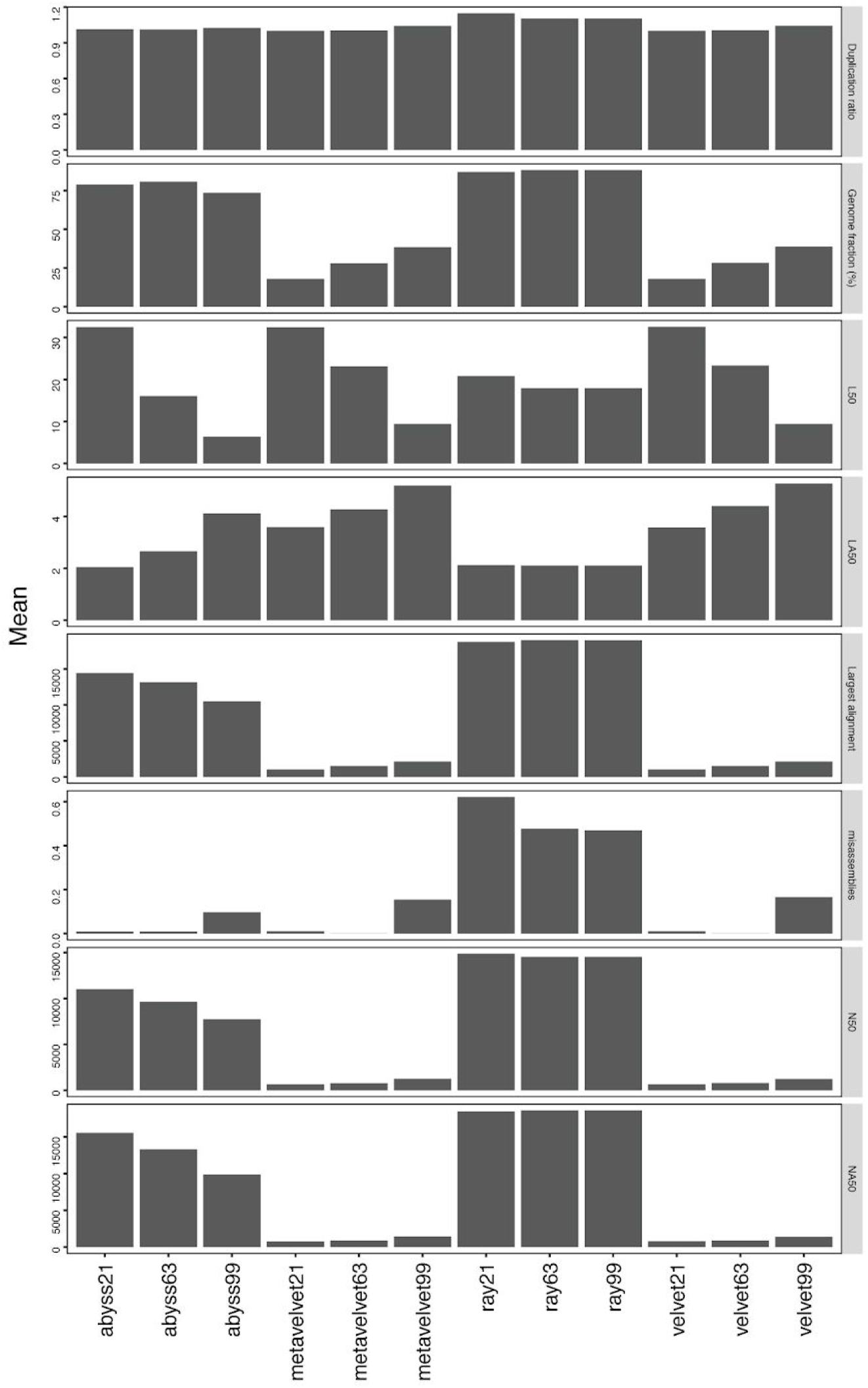
Assembly quality at different k-mers.

**Figure S3.**
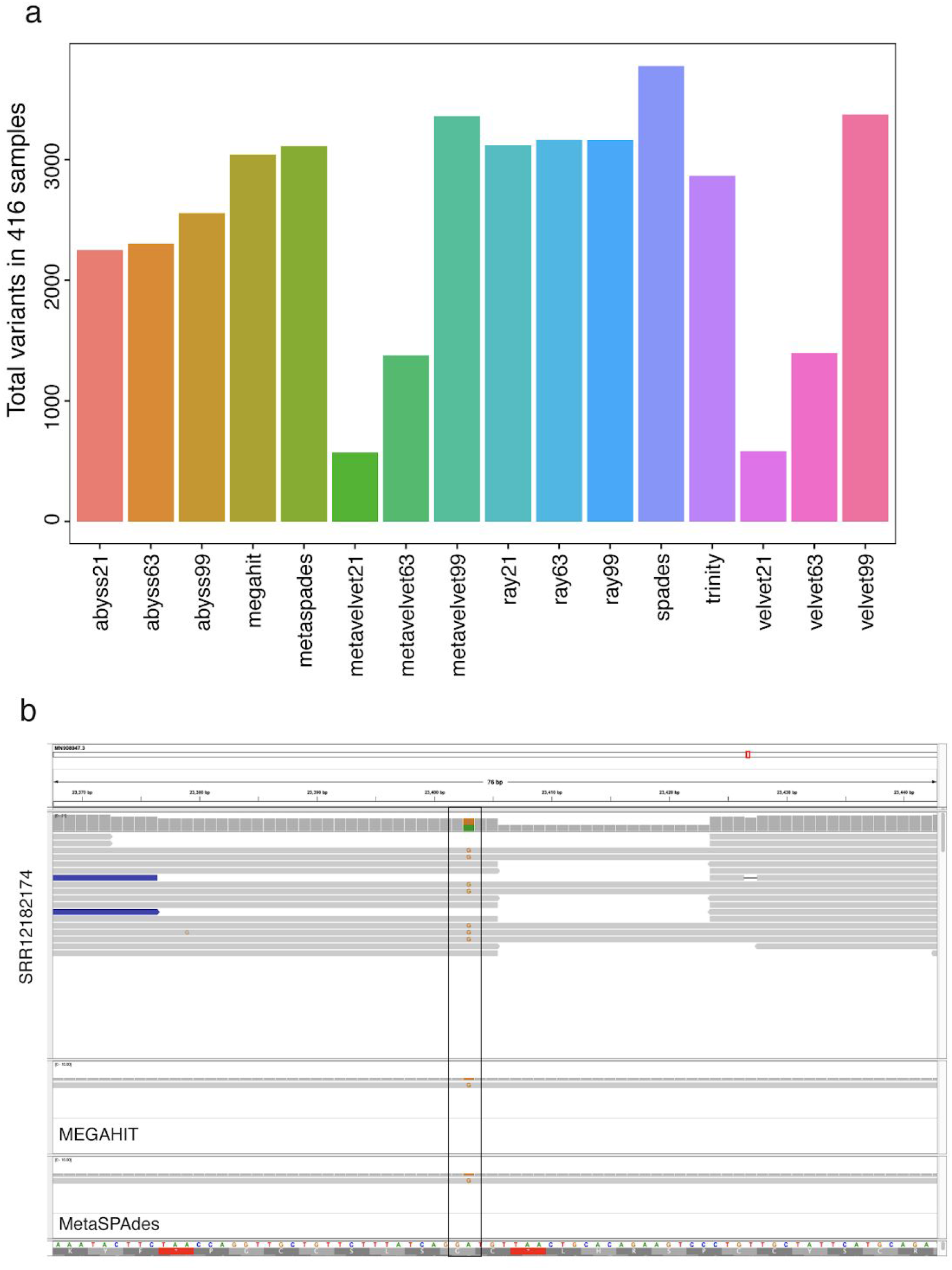
**a)** Variant counts per assembler. **b)** Example of variant presents in both assembly and raw reads.

**Figure S4.**
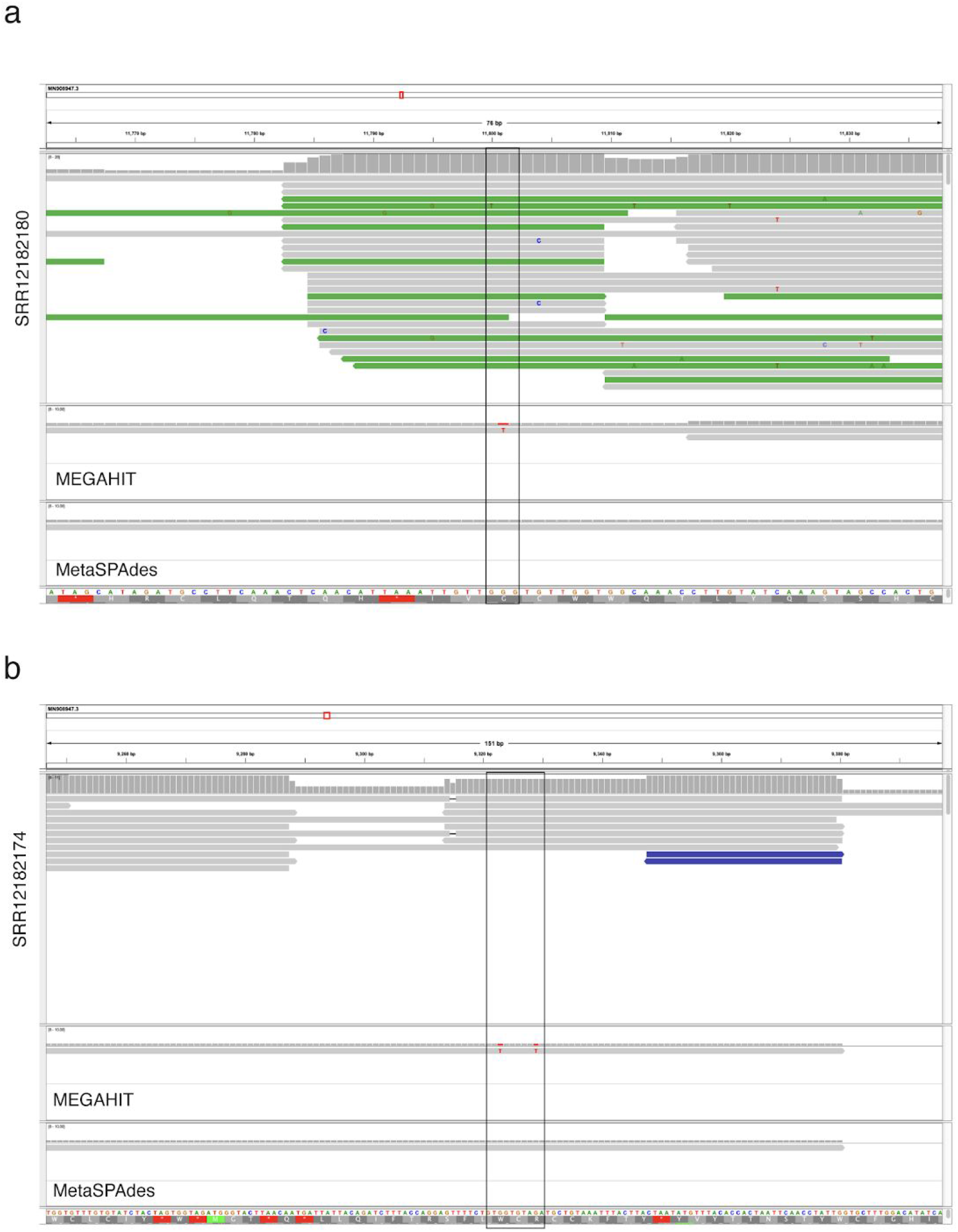
**a,b)** Examples of variants present in one of the assemblies by MEGAHIT and MetaSPAdes but not present in raw reads.

**Table S1.**
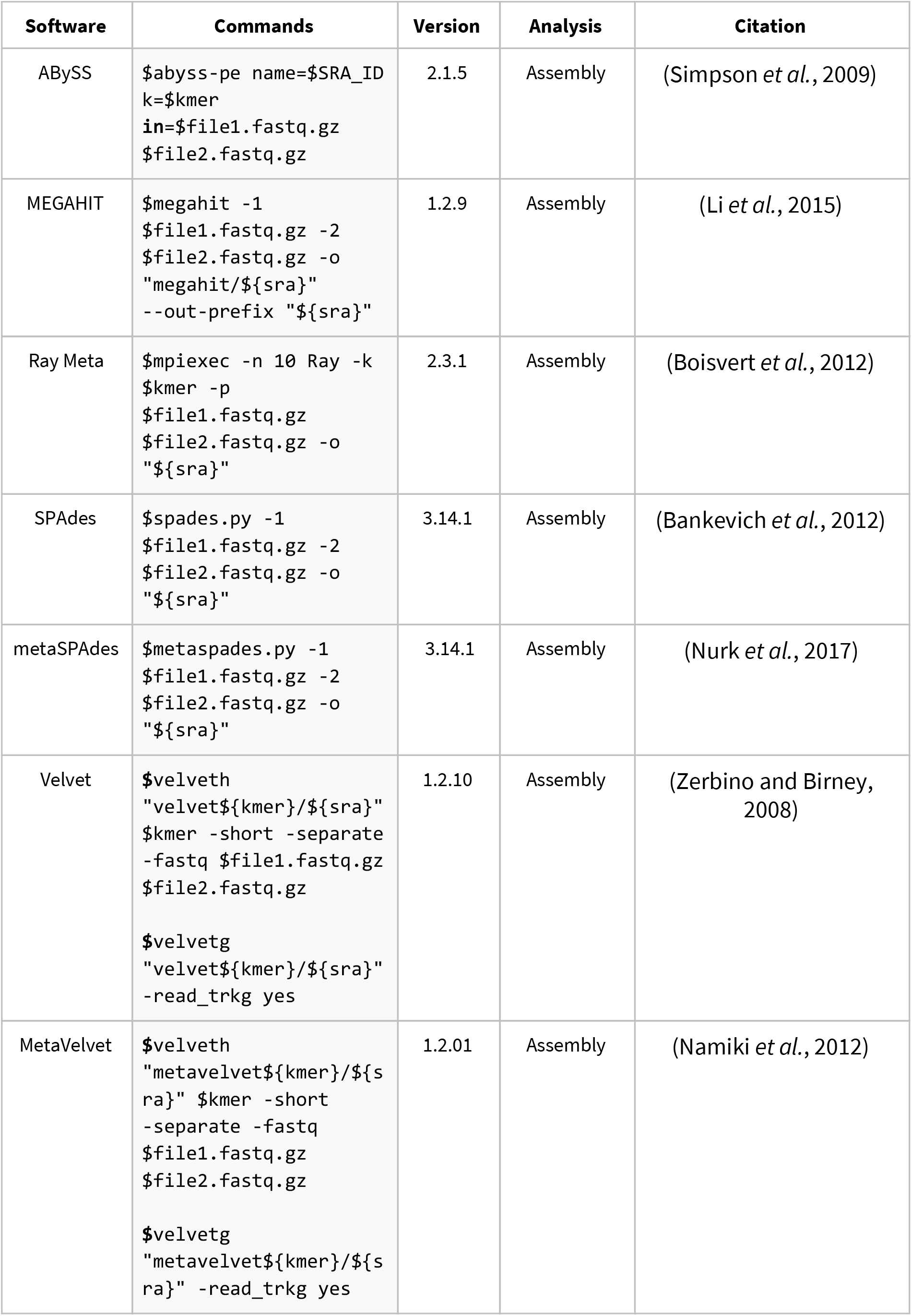
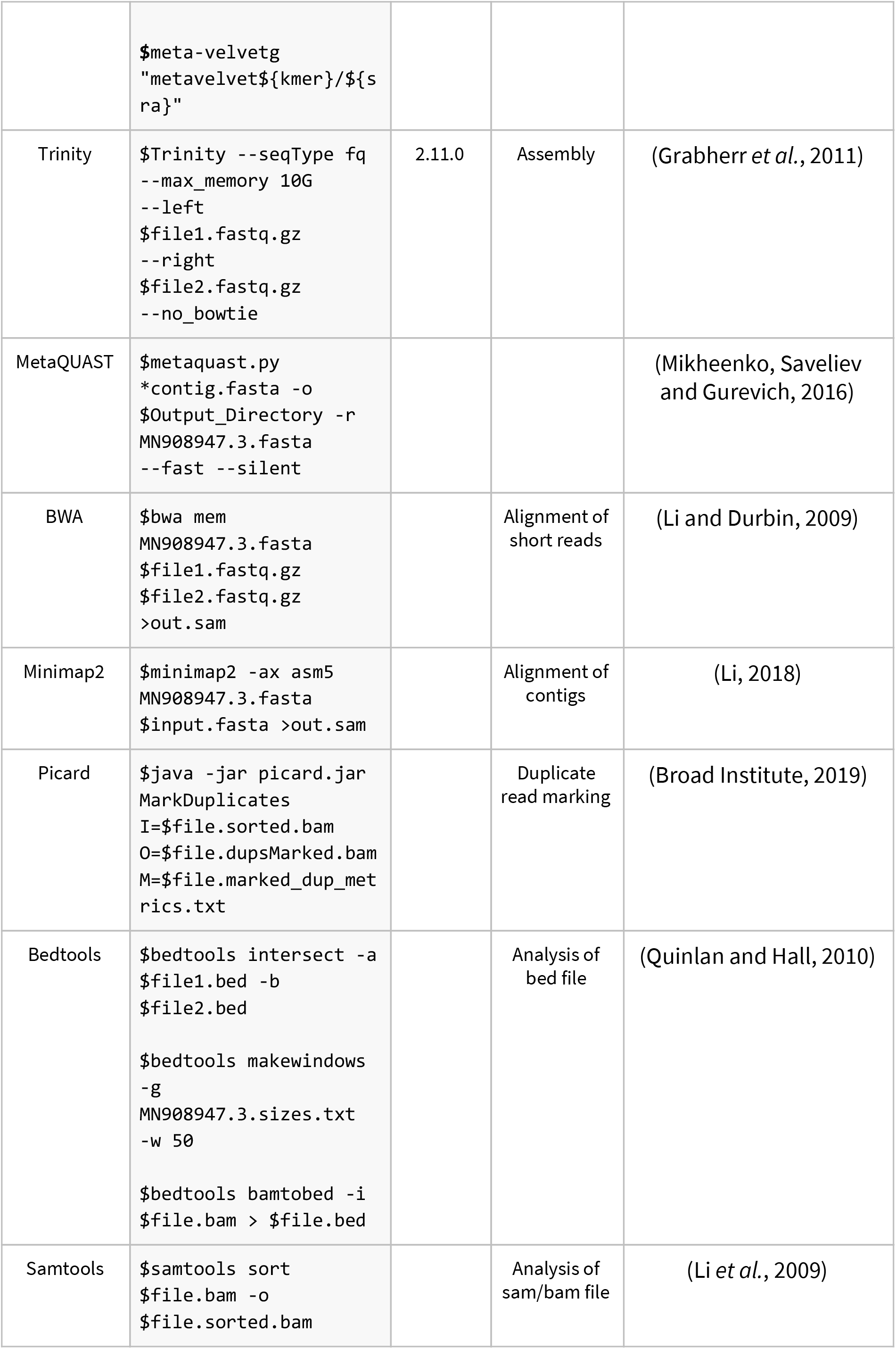
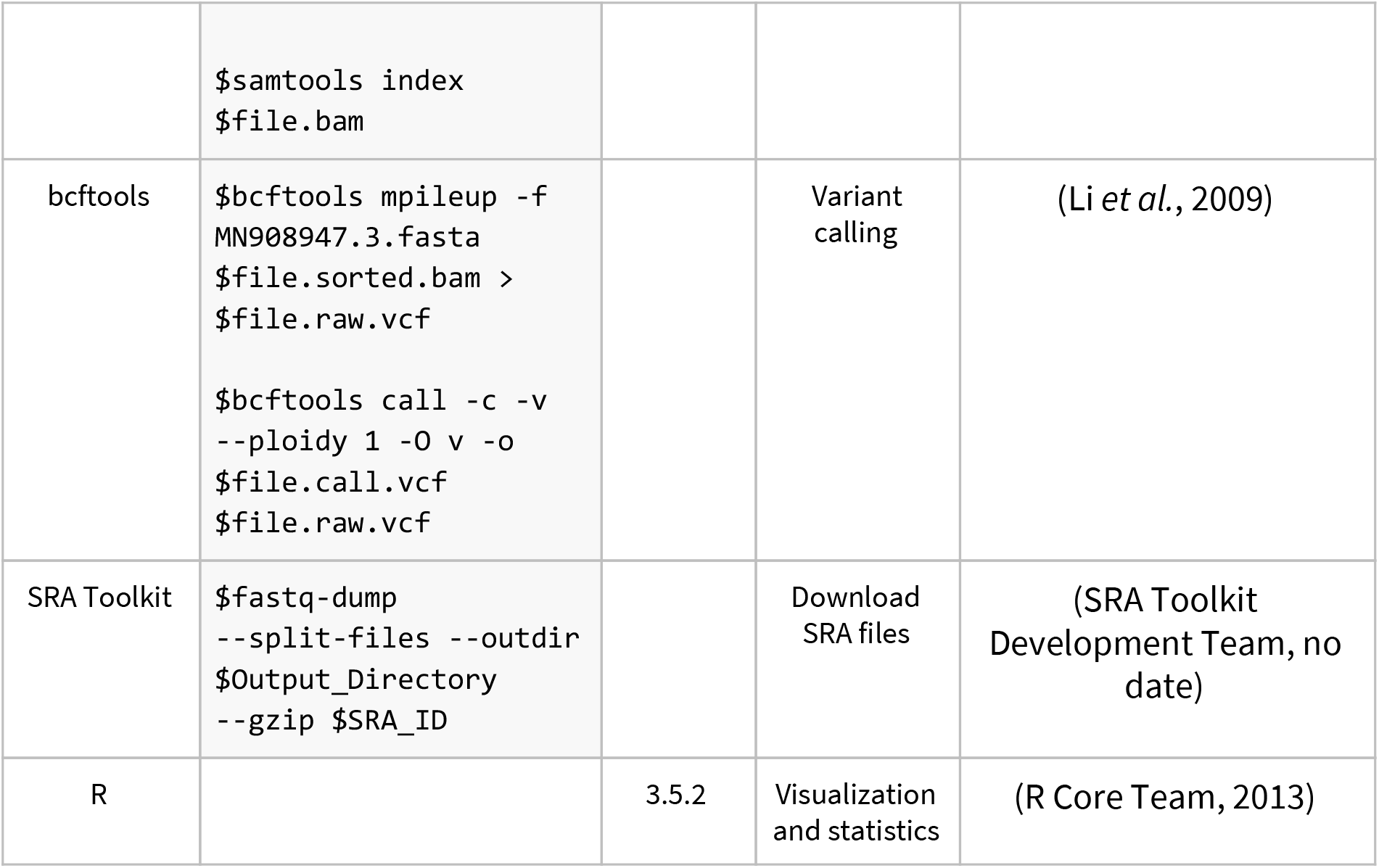
List of software and commands used in this study

**Table S2.** Terminologies used to assess the quality of the genome.

| **Matrices** | **Definition** |
| --- | --- |
| Genome fraction | The ratio of total aligned bases by the assembler to the reference |
| Largest contig | The length of the largest contig in the assembly |
| Nx (N50/N75) | Shortest contig length needed to cover x% of the genome |
| NAx (NA50/NA75) | Combination of the well known Nx metric and Plantagora's number of misassemblies metric. |
| Lx (L50/L75) | The minimum number of contigs that make up x% of genome size |
| LAx (LA50/LA75) | The minimum number of aligned contigs that make up x% of genome |
| Total length | The total number of bases in the assembly |
| Largest alignment | The largest uninterrupted alignment in the assembly |
| Mismatches per 100 kbp | The average number of mismatches per 100 kbp aligned bases |
| Contigs >= 1000 bp | The number of contigs that are >= 1 kb |
| Number of Contigs | The total number of contigs in the assembly |
| Indels per 100 kbp | The average number of single nucleotide insertions or deletions per 100 kbp aligned bases |
| Total aligned length | The total aligned bases of in the assembly |
| Misassemblies | The number of misassemblies |

**Supplementary file1:** Median of assembly quality scores for all samples (Median_Assembly_Quality_assemblers.csv)

**Supplementary file2:** Assembly quality scores for all samples (report_metaquast_PE_all.tsv)

